# Computational redesign of a thermostable T7 RNA polymerase

**DOI:** 10.1101/2025.11.12.688101

**Authors:** Zachary T. Baumer, Timothy A. Whitehead

## Abstract

T7 RNA polymerase (T7 RNAP) is a foundational enzyme for biotechnology, but its utility for many potential applications is limited by low thermal stability of 43-44°C. While stabilized variants exist, the most stable commercial version has a proprietary sequence. In this work we developed a highly stable T7 RNAP using structure-based computational design. We combined mutations from previous stabilized variants (M5, M8, V7abcd) with new mutations identified by PROSS. These mutations were filtered using data-driven heuristics to preserve function. Our final design, T7^T+^, contains 30 point mutations from the original T7 RNAP and demonstrates a functional stability (T_50_) of 54.9°C in a thermal challenge assay, which is 2.4°C higher than the most stable, published open-source variant to date. Circular dichroism spectroscopy showed an apparent melting temperature of 53.8°C. T7^T+^ retains 59% of wild-type activity at 37°C. 16 of the 18 tested protein designs had higher stability against thermal challenge compared with the genetic background, attesting to the high success rates of existing non deep learning computational methods for the design of stable, functional proteins. A plasmid encoding T7^T+^ has been deposited in AddGene and is freely available for non-commercial use.

## Introduction

The single subunit T7 RNA polymerase (T7 RNAP) is a foundational enzyme for biotechnology. Native and engineered versions of T7 RNAP are widely used for instantiation of synthetic biology gene circuits (Baumer *et al*., 2024; McSweeney *et al*., 2025), for recombinant expression of proteins in bacteria(Studier, 2005), and for *in vitro* transcription of therapeutic mRNA (Dousis *et al*., 2023). Thermally stabilizing T7 RNAP can expand its utility in biotech. *In vitro* transcripts produced at higher temperatures have reduced dsRNA formation, which lessens immunogenicity (Wu *et al*., 2020). Further improvements to *in vitro* transcription processes may be realized using immobilized thermostable enzymes to improve downstream purification(Sheldon *et al*., 2021), enable continuous manufacturing (Thompson *et al*., 2019), and increase reusability of the immobilized enzymes (Maghraby *et al*., 2023). Additionally, a thermostable T7 RNAP could be applied in thermophilic organisms, where microbes grow at temperatures in excess of 55 °C. Industrial use of thermophiles has comparative advantages in sterility, available metabolic pathways and feedstocks, and cooling costs (Vavitsas *et al*., 2022).

Native T7 RNAP has an apparent melting and inactivation temperature of 43-44°C at near-neutral pH. Earlier efforts to stabilize T7 RNAP used thermally challenged survival-based screens to identify mutations, which when combined resulted in the M5, M8, and V7abcd variants(Ikeda *et al*., 1992; Sugiyama *et al*., 2009; Boulain *et al*., 2013). The best variants with fully disclosed sequences have thermal stabilities of approximately 52 °C (Boulain *et al*., 2013). A recent paper used PROSS 2.0 (Weinstein *et al*., 2021) and mutational combinations to stabilize a T7 RNAP variant with a melting temperature below 50 °C (Liu *et al*., 2025). Recent deep learning approaches have identified stabilizing mutations (Jiang *et al*., 2024, 2025), but the published variants do not approach the thermal stabilities of the earlier described proteins. New England Biolabs scientists used a computational approach to model mutations which were predicted to be stabilizing using the BioLuminate® software package from Schrodinger (Ong *et* al., 2019). Combining stabilizing mutations led to the highly stabilized Hi-T7® with activity up to 56°C. The specific sequence has not been disclosed, which motivates development of an open source, thermally stable T7 RNAP. In this contribution we used structure-based computational design to further engineer stability into T7 RNAP. We verify stability using *in vitro* thermal challenge activity assays and biochemically characterize the best candidate. Our enzyme, T7^T+^, has the highest stability against thermal challenge (T_50_ 54.9 °C) of published sequences to date and has 59% of the relative activity of WT T7 RNAP at 37°C.

## Methods

### Protein generation, expression, and purification

We constructed all T7 designs using the base plasmid pT7-911Q-His6-WT(Rio, 2013), which was a kind gift from the B. Seelig lab at the University of Minnesota. pT7-911Q-His6-WT encodes for an N-terminally His-tagged (MRGSHHHHHHGS-) T7 RNAP K387R expressed from a pT5-lac promoter. K378R existed in the original gene obtained from the Seelig lab, but is conserved in multiple sequence alignments and appears to not have any noticeable effects. M5 was generated by multisite nicking mutagenesis (Kirby *et al*., 2021) using pT7-911Q-His6-WT as a template. Other plasmids were constructed from gene fragments using Gibson Assembly (Gibson *et al*., 2009). All primers, gBlocks, and plasmid sequences are listed in the SOURCE_DATA_FILE.xlsx available from Zenodo (doi: 10.5281/zenodo.17593694). All gene sequences were confirmed using Sanger sequencing. Proteins were expressed in BL21(DE3) *E. coli* and purified (estimated >95%) by standard Ni-IMAC as described previously (Baumer *et al*., 2024). An example SDS-PAGE gel is shown in **SOURCE_DATA_FILE.xlsx**. Protein concentration was estimated using the Edelhoch method with calculated extinction coefficients(Edelhoch, 1967).

### Computational and rational design

All fragments were constructed in the M5/K387R background (K378R, S430P, N433T, S633P, F849I, F880Y). To identify potential stabilizing mutations, five structures (PDB IDs: 1MSW, 1CEZ, 1QLN, 3E2E, 1H38)(Cheetham and Steitz, 1999; Cheetham *et al*., 1999; Tahirov *et al*., 2002; Yin and Steitz, 2002; Durniak *et al*., 2008) of T7 RNAP were input to PROSS (Goldenzweig *et al*., 2016) using the default settings. For each PDB file, separate runs were performed for each chain of multimeric structures (e.g. 1H38-A, 1H38-B, 1H38-C, and 1H38-D). Each run was also performed using both the Talaris and ref2015 energy functions, resulting in a total number of 144 designs (16 jobs; 9 designs per job). Mutations were selected by compiling and ranking the number of times each position and mutation were predicted to be stabilizing. Structural analysis of high ranking mutations were performed manually, selecting mutations which were surface exposed, Cɑ least 8 Å from the Cɑ of other mutated positions, and distant from the specificity loop (Cheetham *et al*., 1999) and domains important to transcriptional activity (Bonner *et al*., 1992). The total mutations considered was reduced to 149 mutations through ranking (**PROSS_outputs.xlsx** available on Zenodo) before being manually curated to the final fragments and mutations listed in **Table 1**.

**Table 1.**
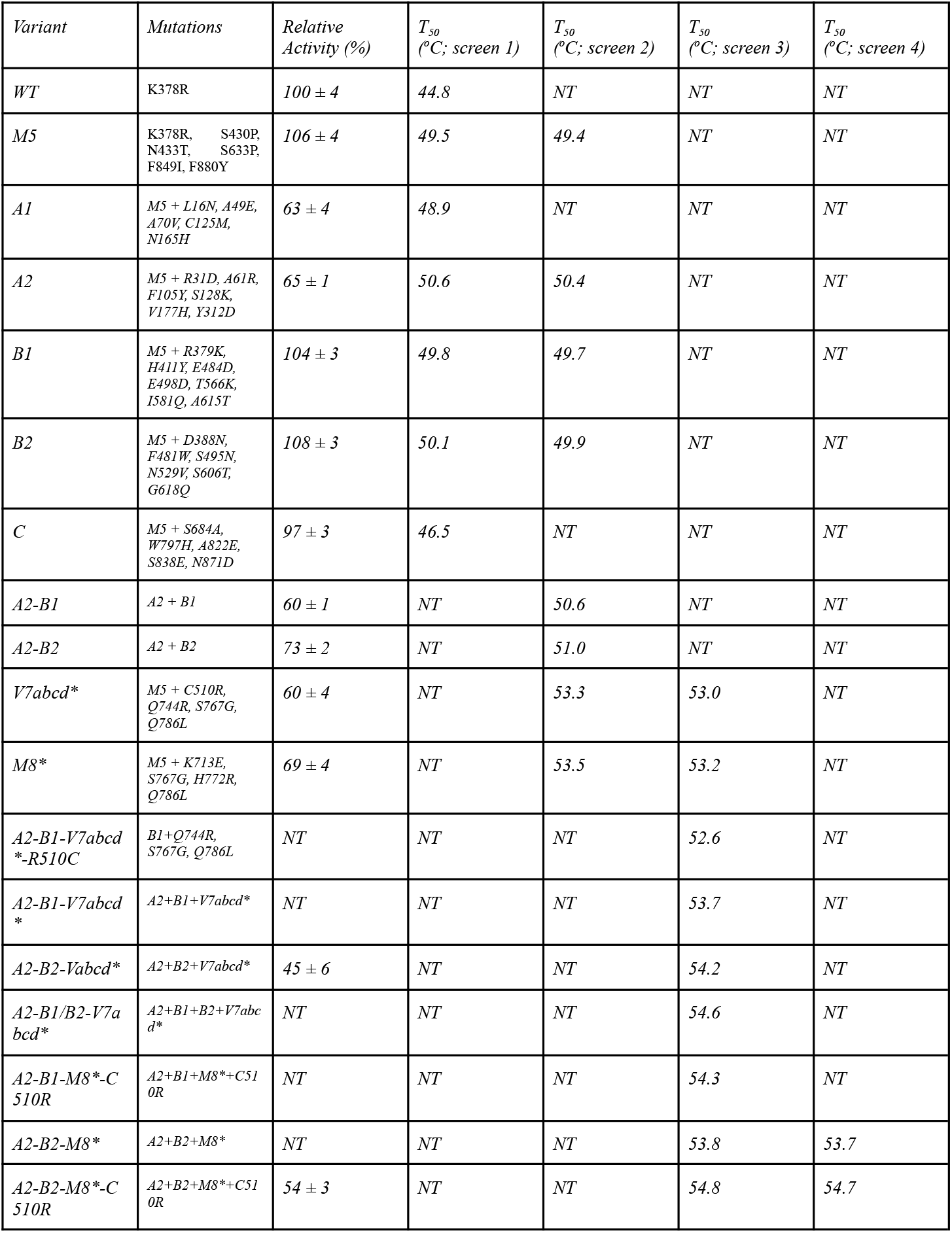
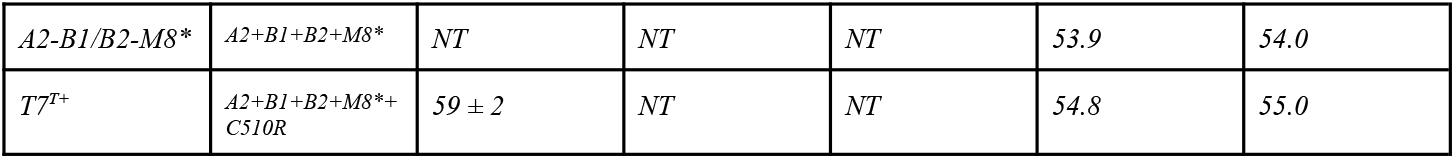
Screening of T7 RNAP designs. Screens represent individual replicates for indicated variants. NT: not tested. Error bars represent 1 s.d. (n ranges from 3 to 9). Relative activity is defined as the activity of a given variant relative to WT.

### Biophysical assays

Thermal challenge assays were performed to measure the functional stability of T7 RNAP. 10 μM of protein was heated in storage solution (50mM Tris-HCl pH 7.9, 1mM EDTA, 5mM beta mercaptoethanol, 100mM NaCl, 15 (v/v) % glycerol) for 25 minutes in a thermocycler set with a temperature gradient (35-60 °C), then brought back to 4°C for at least 20 min. Thermally treated protein was then used in an *in vitro* transcriptional assay at 37 °C as previously described (Baumer *et al*., 2024). Transcription of the Spinach aptamer (Paige *et al*., 2011) sequence yields fluorescence with addition of the ligand 3,5-difluor-4-hydroxybenzylidine imidizolanone. Activity is defined as the maximum rate of fluorescence increase as measured over 20 minute intervals [RFU/minute] which occurred in the first 30 minutes of the reaction start. For thermal challenge assays, the apparent T_50_ was calculated as the temperature at which activity drops to 50% of the maximum measured activity for each variant. The temperature for this analysis was the setpoint temperature for a row of wells in the thermocycler. The T_50_ was derived by fitting a modified power law to the normalized data in GraphPad PRISM.

Circular dichroism spectroscopy was performed on an Applied Photophysics Chirascan V100. Purified protein was buffer exchanged into 20 mM NaPO4, pH 7.8 using 7 kDa Zeba 0.5 mL spin columns per manufacturer’s protocol and diluted to a concentration in the range of 0.1-0.4 mg/mL. Spectral measurements were taken using 1 sec timepoints at 2 nm bandwidth and 0.5 nm step size. For CD melt curves, the wavelength for measuring ellipticity was maintained at 222 nm and the ellipticity was measured for 24 secs between 0.5 °C temperature steps as the sample was heated at 1 °C/min. The actual temperature of the sample was monitored and used for the analysis of the T_m,app_. Instead of fitting data to a two state model, we evaluate the T_m,app_ as being the temperature where the maximum derivative of ellipticity as a function of temperature occurs. We wrote a custom Python script to evaluate T_m,app_. First, a Gaussian filter was applied to smooth the data (gaussian_filter1d, sigma=1). T_m,app_ was taken as the maximum of the derivative evaluated by “numpy.gradient”.

## Results

We used the computational design server PROSS 2.0 (Weinstein *et al*., 2021) to identify potentially stabilizing mutations for T7 RNAP. The stabilizing effect of individual mutations suggested by PROSS are, on average, small (Tsuboyama *et al*., 2023). Thus, many potential mutations need to be combined to see a potent stabilizing effect. However, there is a stability-activity tradeoff with proteins, where increases to stability often lead to decreases in activity, and vice-versa (Stimple *et al*., 2020). PROSS combines individually stabilizing mutations using Rosetta repacking, which mainly serves to prevent spatially proximate mutations. Given the large size of T7 RNAP (883 residues) and its complicated transitions between the initiation and elongation complex, we hypothesized that stacking many potential stabilizing mutations using default PROSS would be ultimately detrimental to function. Therefore, we combined PROSS suggested mutations using a more conservative approach. First, we performed PROSS runs on all available T7 RNAP experimentally determined structures. PROSS predicted 657 stabilizing mutations over 365 of the 883 positions. Second, we filtered these PROSS outputs by the number of times a mutation was observed, as well as selecting only positions that are (i.) distal from active sites, with (ii.) low contact numbers, that involve (iii.) mutations are evolutionary conserved, and (iv.) do not involve mutations to and from proline. We have found that filtering mutations by these relatively crude heuristics (Klesmith *et al*., 2017) improves the redesign success for functional, stabilized proteins (Wrenbeck *et al*., 2019; Beltrán *et al*., 2022; Daffern *et al*., 2024). Third, we tested 5-7 stabilizing mutations at a time in a gene fragment corresponding to a portion of the protein (**Figure 1A-B**) and in the M5 background. Gene fragments containing stabilizing mutations were then recombined into the final design.

**Figure 1.**
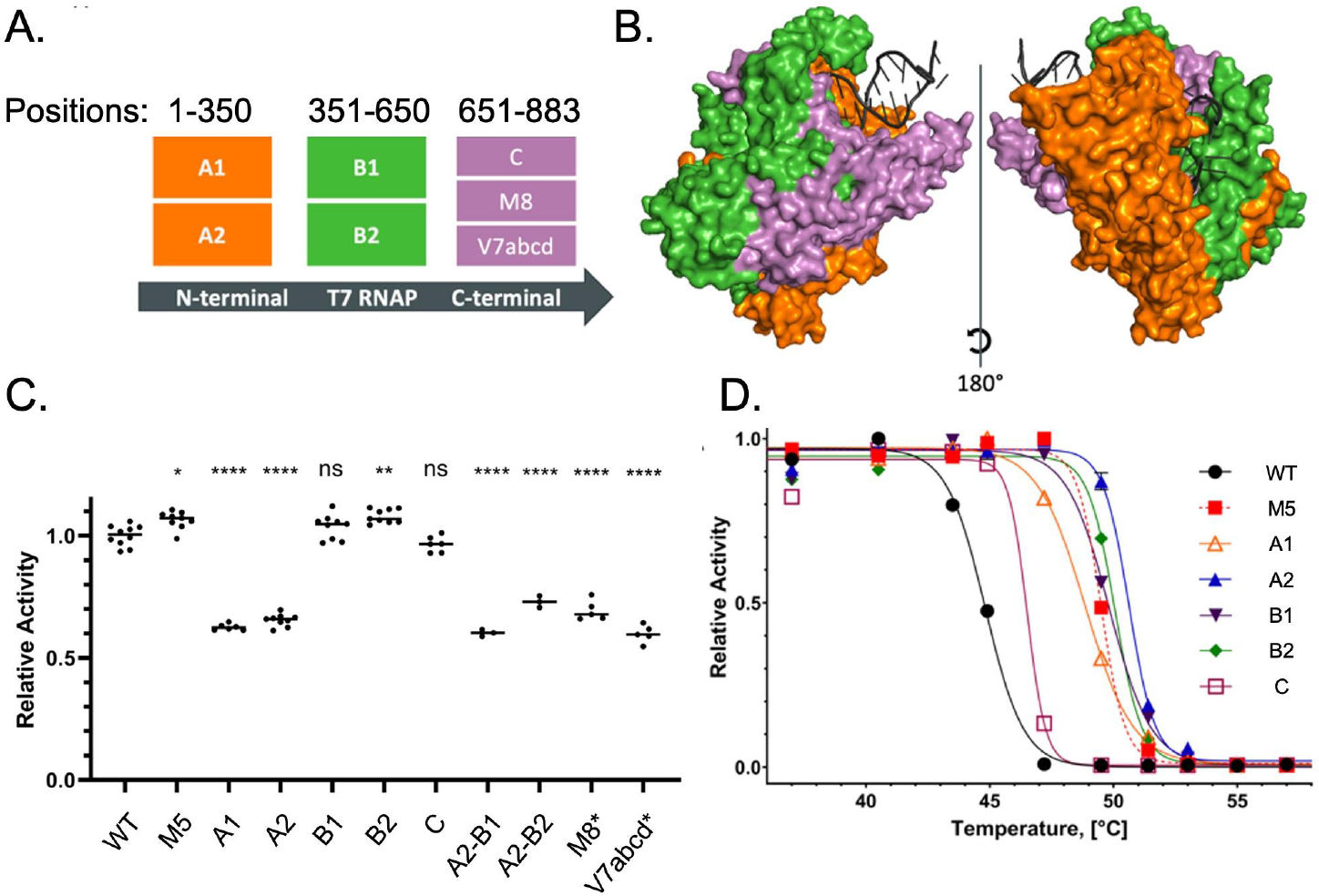
Fragment-based computational design of thermostable T7 RNAP. **A**. Division of the gene encoding T7 RNAP into fragments. Fragment boundaries listed are shown by amino acid positions. **B**. Fragmented T7 RNAP depicted by corresponding colors shown on the initiation complex of T7 RNAP (PDB ID: 1MSW). **C**. Relative activity of T7 RNAP designs compared with WT. Except for WT, all enzymes tested were in the M5 genetic background. p-values were determined using Brown-Forsyth and Welch’s ANOVA, corrected for multiple comparisons. *, <.05; **, <.01; ****, <1e-4; ns, not significant. **D**. Stability against thermal challenge for indicated variants. T7 RNAP designs are incubated at the indicated temperature for 25 minutes before a temperature ramp to 4°C for 20 min. Proteins are then diluted and assayed for activity at 37°C as in panel C. A T_50_ is calculated as the temperature with 50% relative activity of the starting protein. Screen 1 is shown; information on all screens is shown in **Table 1**.

In our initial screen, we tested the relative activity and stability after thermal challenge (T_50_) of five designs using a previously described fluorescence aptamer assay (Baumer *et al*., 2024; Delaney *et al*., 2024). We found that two of five designs (A1, A2) led to statistically significant decreases in relative activity (**Figure 1C, Table 1**) to 63-65% of the WT activity (p value <1e-4,, Brown-Forsyth and Welch’s ANOVA, corrected for multiple comparisons). Interestingly, the B2 design led to a modest but statistically significant increase in relative activity (**Figure 1C**) (p value <0.01). Three of the five designs (A2, B1, B2) showed increases in T_50_ compared with the M5 background (**Figure 1D, Table 1)**. However, these stability increases were modest (ΔT_50_ ranging from 0.3-1.1°C) relative to the M5 background. Our one design incorporating mutations on the C-terminal domain of T7 RNAP (C; positions 651-883) was destabilizing. Thus, in a second round of screening we tested addition of the M8 and the V7abcd mutations in the M5 background (Boulain *et al*., 2013) (M8*, V7abcd*, respectively). These stabilizing mutations target the C-terminal domain of T7 RNAP, which has lower stability than the rest of the protein (Protasevich *et al*., 1994; Meyer *et al*., 2015). The M8* and V7abcd* were considerably more stable than the PROSS designs but at the expense of lower relative activity compared to WT (**Table 1**). For the third screening round, we assessed the T_50_ of nine different combinations of A1, B1, B2, M8, and the V7abcd mutations. We also assessed the relative activity for designs with the highest T_50_. This initial screen of 18 proteins resulted in a lead candidate T7^T+^ (R31D, A61R, F105Y, S128K, V177H, Y312D, K378R, R379K, D388N, H411Y, S430P, N433T, F481W, E484D, S495N, E498D, C510R, N529V, T566K, I581Q, A615T, S606T, G618Q, S633P, K713E, S767G, H772R, Q786L, F849I, F880Y) which had the highest T_50_ and comparatively highest relative activity. T7^T+^ contains 30 total mutations from WT T7 RNAP sourced from three PROSS-designed gene fragments (A2, B1, B2; this work) and three previous stabilized T7 RNAPs (Sugiyama *et al*., 2009; Boulain *et al*., 2013).

We prepared fresh protein preps (SOURCE_DATA_FILE.xlsx) and then assessed T7^T+^ compared with WT T7 RNAP, the M5 variant, and M8*. Our results are presented in **Figure 2**. T7^T+^ has a relative activity of 59%, which is similar to that of the M8* variant. We used circular dichroism (CD) spectroscopy to examine the secondary structure and to test the apparent melting temperatures (T_m,app_) of the four T7 RNAPs. All four enzymes showed essentially identical CD spectra (**Fig 2**). A previous study of the thermal melting and urea unfolding of T7 RNAP by CD (Griko *et al*., 2001) showed that wild-type T7 RNAP undergoes three distinct transitions during its unfolding, likely caused by separate domains unfolding. Thus, a two-state unfolding model is not appropriate to estimate the melting temperature of T7 RNAP; instead, we estimate the T_m,app_ by the midpoint of the molar ellipticity at 222 nm. Using this definition, WT has a T_m,app_ = 43.2°C, consistent with previous measurements. T7^T+^ has a T_m,app_ = 53.8°C, which is >2°C of any other variant. Interestingly, as we inspect the CD-melt curves for T7 RNAP, a steeper transition arises with increasing stability for the M8* and T7^T+^ designs, which may be interpreted as the individual unfolding transitions becoming closer in temperature. We also assessed T_50_ using the previously described thermal challenge assay. T7^T+^ has a T_50_ of 54.9°C, which is 2.4°C higher than any other variant.

**Figure 2.**
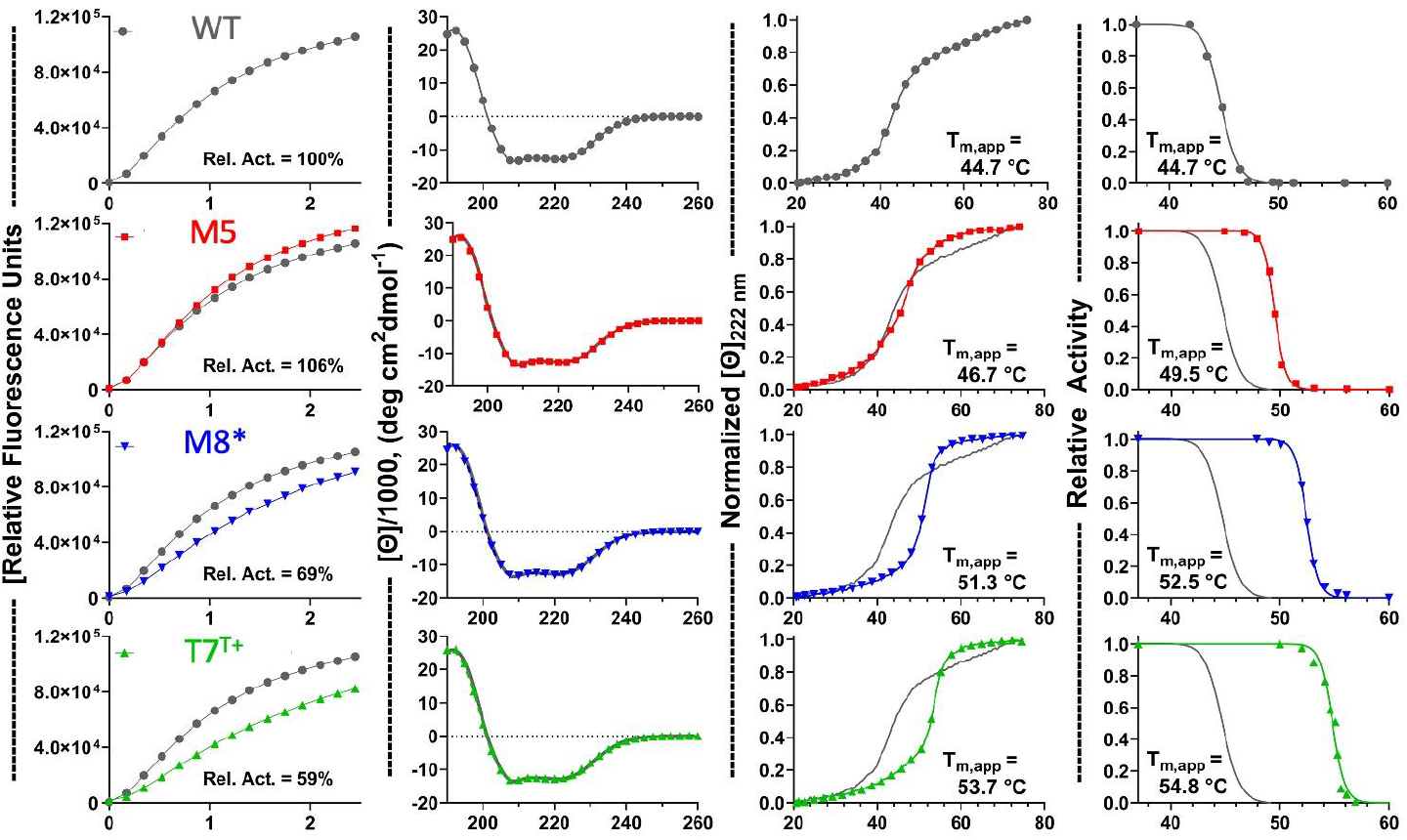
Comparison of activity and apparent Tm of differentially stabilized T7 RNAPs. **A**. Wild-type T7 RNAP; **B**. M5; **C**. M8*; **D**. T7^T+^; **[Column I]** Activity; **[Column II]** Circular Dichroism near-UV spectra; **[Column III]** Temperature melts using circular dichroism spectroscopy. Molar ellipticity was measured at 222nm during the temperature ramp; **[Column IV]** Thermal challenge activity assay, normalized to activity at 37 °C. Activity plots are the mean trace of two separate runs with the listed “relative activity”. CD Spectra are given as a single spectra that was averaged from two traces during measurement. CD Melt curves are given as a single melt curve for each variant, except T7^T+^ which was performed in replicate. Thermal challenge activity assays are the average of two technical replicates.

## Discussion and Conclusion

Estimates from deep mutational scanning suggest that on the order of 1% of random mutations are stabilizing without impacting enzyme function (Klesmith *et al*., 2017). For T7 RNAP, this means that there are approximately 150 thermally stabilizing point mutations that could be recombined into stable variants. Our design, T7^T+^, contains up to 30 of these mutations, including 19 identified in the present work. Not all may be stabilizing, and further benefit may be found by testing these and other PROSS-suggested mutations individually. For example, the stabilizing effects of C510R first identified in the Vabcd variant were evident in all the constructs tested (**Table 1**). Further, there is minimal overlap between mutations in our design and recent thermostabilized T7 RNAPs. While the NEB patent (Ong *et al*., 2019) does not disclose the exact sequence, it does list potential mutations; 3 out of 19 mutations are shared. Taken together, the body of evidence suggests that there is further room for improvement in T7 RNAP with regards to thermal stability.

While deep learning is in vogue across science, our results show that existing computational methods grounded in biophysical principles still have considerable utility. We were able to identify T7^T+^ by screening only 18 proteins in total; of these, 16 were more thermally stable than the M5 genetic background used in the experiment. We did not demonstrate further application of T7^T+^ as it is outside the scope of this short communication. To expedite adoption of T7^T+^ in the broader community, we have deposited a plasmid expressing T7^T+^ in the AddGene plasmid repository (Kamens, 2014). This construct is freely available for non-commercial use.

## Data and Code Availability

All data and novel code used in this paper is available in a Zenodo repository (doi: 10.5281/zenodo.17593694). This repository contains:(1.) SOURCE_DATA_FILE.xlsx contains all experimental data presented in this paper. This file also contains all primers, plasmid sequences, and gBlock sequences used for plasmid construction. (2.) The Jupyter notebook ‘Evaluating Tmapp from CD.ipynb’ is a custom script for extracting T_m,app_ from circular dichroism temperature melts. (3.) PROSS_outputs.xlsx lists the full PROSS outputs and our relative ranking of the mutations for testing in this paper. Plasmid pZB246 encoding T7^T+^ is available on AddGene (to be provided upon publication).

## Acknowledgements

This work was supported by the National Science Foundation (Award #: 2218330), and the National Science Foundation Graduate Research Fellowship Program (Z.T.B. DGE Award Number 2040434, fellow ID: 2021324468). We also thank the Shared Instruments Pool (RRID: SCR_018986) of the Department of Biochemistry at the University of Colorado Boulder for the use of the CD spectrometer. The CD is funded by NIH Shared Instrumentation Grant S10RR028036.

